# MicroRNA-focused CRISPR/Cas9 Screen Identifies miR-142 as a Key Regulator of Epstein-Barr Virus Reactivation

**DOI:** 10.1101/2024.01.15.575629

**Authors:** Yan Chen, Rodney P. Kincaid, Kelley Bastin, Devin N. Fachko, Rebecca L. Skalsky

## Abstract

Reactivation from latency plays a significant role in maintaining persistent lifelong Epstein-Barr virus (EBV) infection. Mechanisms governing successful activation and progression of the EBV lytic phase are not fully understood. EBV expresses multiple viral microRNAs (miRNAs) and manipulates several cellular miRNAs to support viral infection. To gain insight into the host miRNAs regulating transitions from EBV latency into the lytic stage, we conducted a CRISPR/Cas9-based screen in EBV+ Burkitt lymphoma (BL) cells using anti-Ig antibodies to crosslink the B cell receptor (BCR) and induce reactivation. Using a gRNA library against >1500 annotated human miRNAs, we identified miR-142 as a key regulator of EBV reactivation. Genetic ablation of miR-142 enhanced levels of immediate early and early lytic gene products in infected BL cells. Ago2-PAR-CLIP experiments with reactivated cells revealed miR-142 targets related to Erk/MAPK signaling, including components directly downstream of the B cell receptor (BCR). Consistent with these findings, disruption of miR-142 enhanced SOS1 levels and Mek phosphorylation in response to surface Ig cross-linking. Effects could be rescued by inhibitors of Mek (cobimetinib) or Raf (dabrafenib). Taken together, these results show that miR-142 functionally regulates SOS1/Ras/Raf/Mek/Erk signaling initiated through the BCR and consequently, restricts EBV entry into the lytic cycle.

## Introduction

Epstein-Barr virus (EBV) is a human gamma-herpesvirus that infects >95% of adults worldwide. Primary infection is normally asymptomatic in young children; however, individuals that acquire EBV as adolescents or adults often develop infectious mononucleosis (1, 2). Several lymphoproliferative disorders, autoimmune diseases including Multiple Sclerosis, and cancers such as Burkitt lymphoma (BL) and nasopharyngeal carcinoma (NPC) are linked to EBV infection (3, 4). Endemic BL, in particular, is an aggressive form of EBV-associated non-Hodgkin’s lymphoma (NHL) and the most common childhood cancer in the equatorial region of sub-Saharan Africa (5). An estimated 2% of all cancer-related deaths are directly attributed to EBV (6). Following initial infection, EBV establishes a life-long infection that is predominantly latent, with only a few essential latent viral genes expressed. Periodic reactivation, which induces lytic viral gene expression and production of viral progeny, is necessary for cell-to-cell spread, host-to-host transmission, and replenishing the latent reservoir of infected cells (7, 8). Mounting evidence suggests that lytic replication is a key contributor to viral oncogenesis (9, 10). Lytic gene products can be detected in EBV-associated tumors (11), increased EBV DNA loads correlate with BL disease progression in children (12), and elevated antibody titers to EBV lytic antigens often precede onset of cancers such as NPC (13, 14). EBV reactivation is also observed following co-infections with other viruses, such as HIV or SARS-CoV2, and recently, has been linked to fatigue symptoms associated with Post-Acute Sequelae of COVID-19 (PASC), also known as long COVID, adding to the complexity of EBV-associated diseases (15, 16, 17, 18).

Molecular processes regulating EBV reactivation are not fully understood. In B cells, EBV reactivation is tightly linked to cellular differentiation programs. Plasma cell transcription factors, such XBP-1 and the master transcriptional regulator Blimp1/PRDM1, can activate promoters of the EBV immediate early (IE) genes Zta and Rta, thereby triggering the lytic cascade (19, 20, 21). *In vitro,* EBV reactivation can be induced by various chemicals such as phorbol esters and calcium ionophores or through more physiological triggers such as cross-linking of surface immunoglobulins to mimic antigen engagement of the B cell receptor (BCR). BCR-mediated signaling activates multiple downstream pathways such as mitogen-activated protein kinase (MAPK), phosphatidylinositol 3-kinase (PI3K), and nuclear factor–kB (NFkB) that direct transcriptional responses, outcomes of B cell differentiation (22, 23), and in EBV-infected cells, stimulate activity of the Zta promoter (24).

In previous studies, we and others have demonstrated that several microRNAs (miRNAs) play important roles in the EBV latent to lytic switch, consequently modulating lytic gene expression and virus production from infected cells (25, 26, 27, 28, 29). miRNAs are small, non-coding RNAs that post-transcriptionally control gene expression and protein abundance by recruiting the RNA-induced silencing complex (RISC) to sequence-specific sites on target mRNAs, predominantly within 3’UTRs. Accumulating evidence demonstrates that miRNAs play critical roles in disease pathogenesis due to their pleiotropic functions in modulating basic cellular processes and can have activity as tumor suppressors or oncogenes. With regards to EBV infection, specific miRNAs such as miR-155, miR-34a, and miR-17 contribute to the maintenance of latency. Knockdown of either miR-155 or miR-34a impairs the growth of latently infected lymphoblastoid cell lines (LCLs) (30, 31) whereas knockdown of miR-17 in EBV-positive BL cells enhances EBV IE gene expression following BCR engagement (25). miR-155 further suppresses bone morphogenetic protein (BMP) signaling events that lead to EBV reactivation through the direct targeting of SMAD1 and SMAD5 (28). In contrast, members of the miR-200 family promote the EBV lytic phase. miR-200b and miR-429 target ZEB1 and ZEB2, which are both transcriptional repressors of the Zta promoter, and consequently, lytic gene expression in epithelial cells increases (27, 29). In B cells, another miR-200 family member, miR-141, targets FOXO3 to support progression of the EBV lytic cycle when cells are stimulated to reactivate (26).

Despite these few examples, the functions for most cellular miRNAs in modulating EBV infection, reactivation, and lytic cycle progression remain to be elucidated. Here, we performed a CRISPR/Cas9 screen against >1500 human miRNAs to identify key regulators of the EBV latent to lytic switch and subsequently, investigated the miRNA targets and targeted pathways linked to this process.

## Results

### Genome-wide CRISPR/Cas9 screen identifies host miRNAs impacting EBV reactivation

To identify host miRNAs that impact EBV latency and reactivation, we performed a miRNA-focused CRISPR/Cas9-based loss-of-function screen (Fig. 1A). An LX-miR guide RNA (gRNA) library consisting of 8,382 individual gRNA constructs that were optimized to target 1,594 annotated human miRNA genes (∼85% of human miRNA genes with ∼4 gRNAs per miRNA) (32) was introduced in MutuI BL cells stably expressing a tet-inducible Cas9 (MutuI-iCas9) (26). This library also contains ∼1000 non-targeting control gRNAs. After selection for stable incorporation of the gRNA library, cells were expanded, then treated with doxycycline for one week to express Cas9 and induce gRNA activity. The EBV lytic cycle was subsequently activated by treating cells with anti-IgM for 48 hours. Cells were fixed, stained with an antibody against the EBV late viral envelope protein gp350, and then FAC-sorted into gp350 positive and negative populations. DNA was isolated from the two populations and gRNA amplicon sequencing libraries were generated. Following sequencing of the gRNA pools for each population, we used the drugZ algorithm, which identifies both synergistic and suppressor interactions in chemogenetic screens (33), to identify candidate miRNAs that modulate lytic reactivation. This analysis revealed a total of 39 significantly scoring miRNAs (adjusted p-value < 0.05). gRNAs for 17 miRNAs were enriched in the gp350- positive population while 22 were enriched in the gp350-negative population (Fig. 1B, S1A). Focusing on positively scoring miRNA genes, we identified miR-142 as a top scoring miRNA (Fig. 1B, S1A). All four gRNAs against the miR-142 locus were significantly enriched in the gp350- positive population, strongly indicating that miR-142 functionally restricts EBV reactivation.

**Figure 1.**
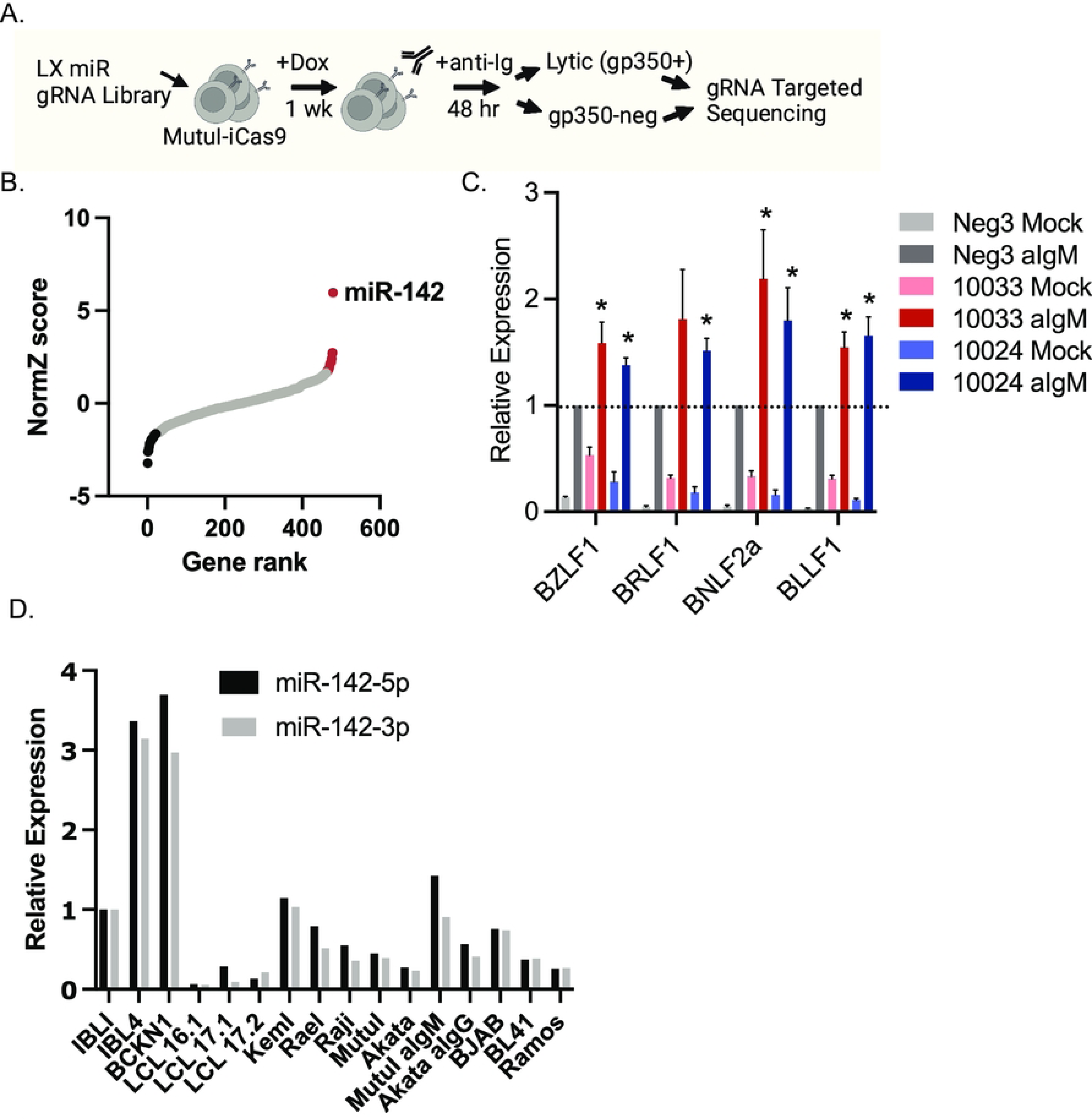
Genome-wide miRNA-focused CRISPR/Cas9 screen identifies cellular miR-142 as a regulator of EBV reactivation. A. Schematic of the miRNA-focused CRISPR/Cas9 loss-of-function screen carried out in EBV-positive MutuI-iCas9 BL cells. B. DrugZ-calculated normZ score plotted versus miRNA gene rank. Red indicates targets that were significantly enriched in gp350-positive population while black indicates targets significantly enriched in the gp350- negative population (p-value < 0.05). Highlighted is miR-142 which showed the most significant changes. C. EBV lytic transcripts are upregulated in miR-142 mutant cells. Four EBV lytic genes were measured by qRT-PCR in MutuI-iCas9 cells transduced with either a control gRNA (Neg3) or the top two scoring guide-RNAs (gRNAs) against miR-142 (10024 or 10033). Cells were either mock or anti-IgM treated for 48 hrs. Values are normalized to GAPDH and shown relative to anti-IgM treated Neg3 control cells. Shown is the average of six independent experiments. By Student’s t-test, *p<0.05. D. miR-142-3p and miR-142-5p expression in multiple B cell lines. Total RNA was extracted from DLBCLs (IBL1, IBL4, BCKN1), LCLs (LCL16.1, 17.1, 17.2), EBV-positive BL (KemI, RaeI, Raji, MutuI, Akata), and EBV-negative BL (BJAB, BL41, Ramos). miRNAs were assessed by Taqman qRT-PCR. Values are normalized to miR-16 and reported relative to IBL1.

To confirm results of the CRISPR/Cas9 screen, we generated individual gRNA vectors targeting the miR-142 locus (gRNA #10033 and #10024). MutuI-iCas9 cells were stably transduced with the two individual miR-142 gRNAs or negative control gRNA (Neg3) and treated with doxycycline for one week to express Cas9 (Fig. S1B and S1C). EBV reactivation was subsequently tested by treating engineered cells with anti-IgM for 48 hrs and monitoring lytic gene expression by qRT-PCR. Compared to negative control cells, MutuI cells receiving miR-142 gRNAs exhibited signs of spontaneous lytic viral transcription in the absence of stimuli, and upon treatment with anti-IgM, demonstrated heightened levels of immediate early, early, and late viral transcripts (Fig. 1C). We observed minimal differences in steady state viral DNA levels in the mock treated cells (Fig. S1B), suggesting that observed phenotypes are linked directly to the transcriptional responses triggered through BCR activation. These findings implicate miR-142 as an important regulator of EBV reactivation in BL cells.

### Expression of miR-142 in transformed B cell lines

Somatic mutations in miR-142 that confer loss of function have been observed in several types of hematological malignancies including DLBCL (diffuse large B cell lymphoma) and AML (acute myelogenous leukemia), and are thought to contribute to oncogenesis by impairing developmental states of hematological cell lineages (34, 35, 36, 37). A point mutation in miR-142 (chr17:58331263C>T[-]) has been described in the EBV-positive Raji BL cell line (35), although it remains unclear whether this impacts miRNA expression and/or function in these cells. An ongoing question is whether EBV infection influences miR-142 levels. miR-142-3p expression was found to be downregulated in EBV-positive compared to EBV-negative NK/T cell lymphoma (38). In contrast, miRNA profiling studies of BL found miR-142-5p to be upregulated in EBV-negative BL cases compared to EBV-positive cases (39). To investigate miRNA levels in EBV- infected B cells, we thus measured miR-142-3p and miR-142-5p in multiple EBV-positive and EBV-negative B cell lines by qRT-PCR (Fig. 1D). miRNA expression in MutuI and Raji cells was comparable to that of other BL cell lines, and anti-Ig treatment only modestly increased both miR- 142-3p and -5p (Fig. 1D). No significant differences in miR-142 levels were found between EBV- negative and EBV-positive BL cells. Strikingly, compared to germinal-center derived BL and DLBCL cell lines, EBV B95-8 LCLs (from three different donors) exhibited the lowest levels of miR-142-3p and -5p (Fig. 1D). Together, these observations support that miR-142 expression is not directly linked to EBV gene products and likely associated with states of B cell differentiation.

### Perturbation of miR-142 enhances EBV lytic cycle

CRISPR/Cas9 targeting of genes in cell lines often results in mixed cell populations with varying degrees of editing. Indeed, assessment of miR-142-5p and miR-142-3p levels within the bulk MutuI-iCas9 10033 and 10024 gRNA populations revealed only modest reductions in miRNA expression (Fig. S1C) despite significant differences in lytic EBV transcripts (Fig. 1C). To maximize targeted disruption of miR-142 in MutuI-iCas9 cells, clonal cell lines were established by limiting dilution. miRNA expression was then assessed by qRT-PCR and two cell lines produced with independent miR-142 gRNAs showing the greatest reduction in miR-142 levels (>60% knockdown) were selected for further characterization (Fig. 2E and 2F). Despite extensive screening, we were unable to achieve clonal cell lines in which miR-142 was fully knocked out, suggesting that complete genetic abrogation of miR-142 from EBV-positive BL might be lethal to these cells. To evaluate miR-142 regulatory functions, we subsequently tested EBV transcripts and expression of lytic proteins in response to reactivation stimuli (Fig. 2A-D). By immunoblot analysis, we detected strong increases in Zta, Ea-D (BMRF1), and BHRF1 protein in both of the miR-142 mutant cell lines (Fig. 2A and B). Consistent with changes at the protein level, expression of immediate early (BZLF1), early (BNLF2a), and late (LMP1) viral genes were also significantly upregulated in miR-142 mutant cells (Fig. 2C and D). Together, these results confirm that targeted disruption of miR-142 alters the course of EBV reactivation and leads to increased production of lytic proteins in response to BCR cross-linking.

**Figure 2.**
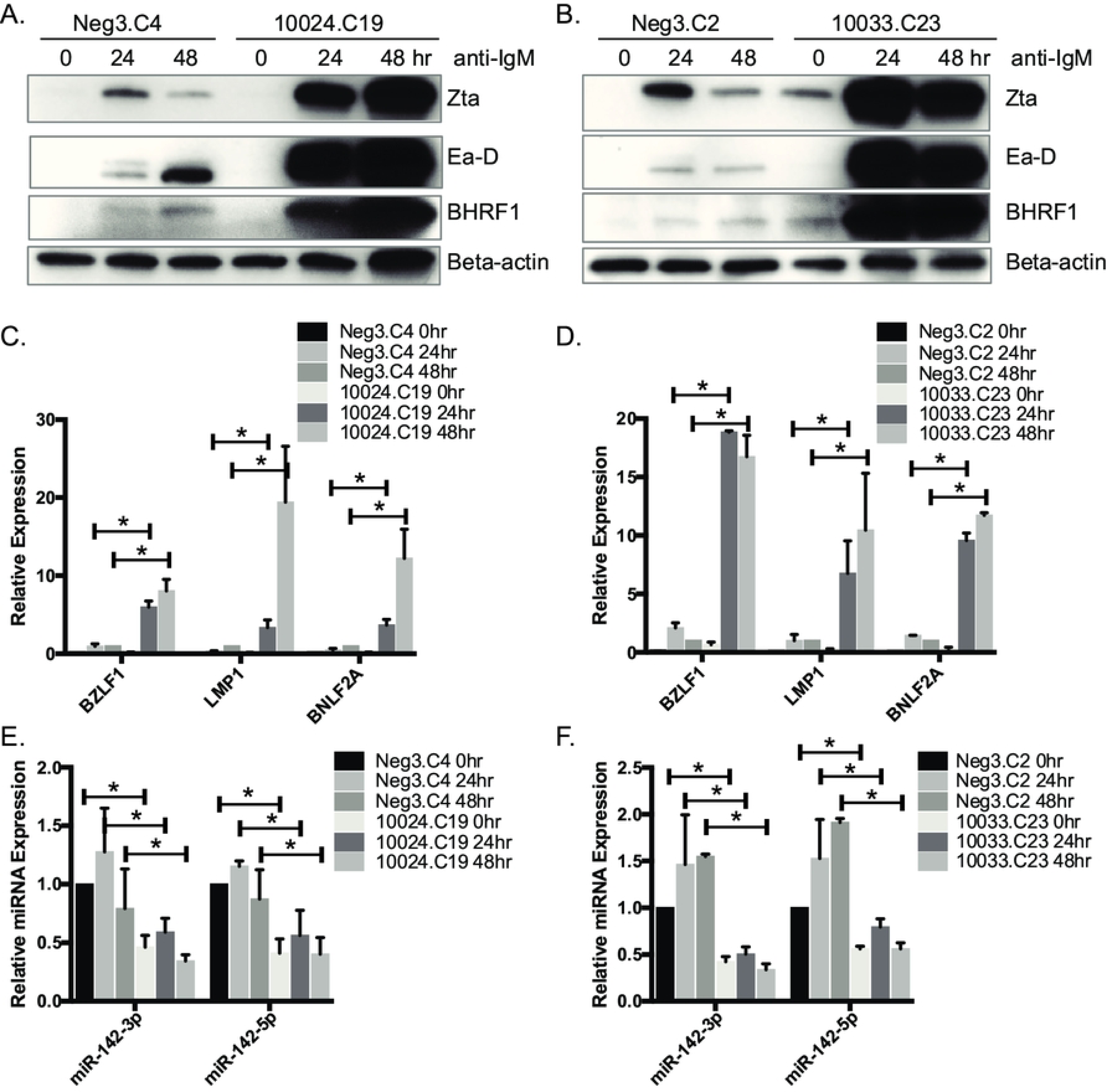
Depletion of miR-142 enhances EBV lytic gene products. A. and B. Immunoblot analysis of EBV viral proteins in Neg3 (C4 or C2 single clones) or miR-142 mutant (10024.C19 or 10033.C23 single clones) BL cells following anti-IgM treatment for 0, 24 or 48 hrs. Beta-actin levels are shown as loading controls. Shown is the representative of three independent experiments. C. and. D. EBV viral gene expression in control (Neg3) or miR-142 mutant cells (10024.C19 or 10033.C23) following anti-IgM treatment for 0, 24 or 48 hrs. Lytic gene expression was determined by qRT-PCR. Values are normalized to GAPDH and shown relative to Neg3 control cells treated with anti-IgM for 48 hrs. Shown is the average of three independent experiments. Student’s t-test, *p<0.05. E. and F. Validation of miR -142 knockdown in miR-142 mutant cells. Taqman qRT-PCR was performed for miR-142-3p or 5p expression in total RNA samples from panels C. and D. Values are normalized to miR-16 and shown relative to Neg3 control cells at 0 hr. Shown is the average of three independent experiments. Student’s t-test, *p<0.05.

### Genome-wide identification of miR-142 targets by Ago-PAR-CLIP

miR-142 is a unique miRNA in that, in addition to canonical 5p and 3p mature miRNAs, natural 5’ isomiRs (miR-142-3p-1 and miR-142-5p-1) are generated from differential Drosha processing of the primary miRNA transcript (40). The major 5’ isomiR, miR-142-3p-1, is shorter than miR-142- 3p at the 5’ terminus by a single nucleotide and thus, harbors a different “seed” sequence and expands the range of sequence-specific targets for these miRNAs (41). To experimentally identify candidate genes that are directly targeted by miR-142 miRNAs or any of the isomiRs, we performed Ago-PAR-CLIP (photoactivatable ribonucleoside-enhanced crosslinking and immunoprecipitation) analysis on MutuI and Akata BL cells. Briefly, mock or anti-Ig treated cells were cultured in the presence of 4-thiouridine to label nascent RNAs, Ago-interacting RNAs were UV crosslinked at 365 nm, and following immunopurification using antibodies against Ago proteins, RNAs were converted into cDNA and sequenced. We generated five PAR-CLIP libraries for MutuI cells and four libraries for Akata cells. A sixth PAR-CLIP library of reactivated MutuI cells is previously published (PRJNA773484) (42) and was included in our downstream bioinformatics analysis. Reads were aligned to the human genome and subsequently analyzed by PARalyzer (43) and microMUMMIE (44) to identify Ago/miRNA binding sites with T-to-C conversions. We focused our analysis specifically on miR-142 interaction sites within 3’UTRs and identified sites in transcripts from >400 unique genes that harbored canonical seed matches (i.e. >=7mer1A) to the miR-142 miRNAs (Table S1). Comparison of the 318 genes identified in mock-treated BL cells with the 232 genes identified in reactivated BL cells revealed 147 targets that were common to both conditions and 85 3’UTRs specifically targeted by miR-142 miRNAs in reactivated cells (Fig. 3A, Table S1). 3’UTR target sites for 187 different genes were detected in at least two or more of the PAR-CLIP libraries (irrespective of condition) and are reported in Fig. 3B.

**Figure 3.**
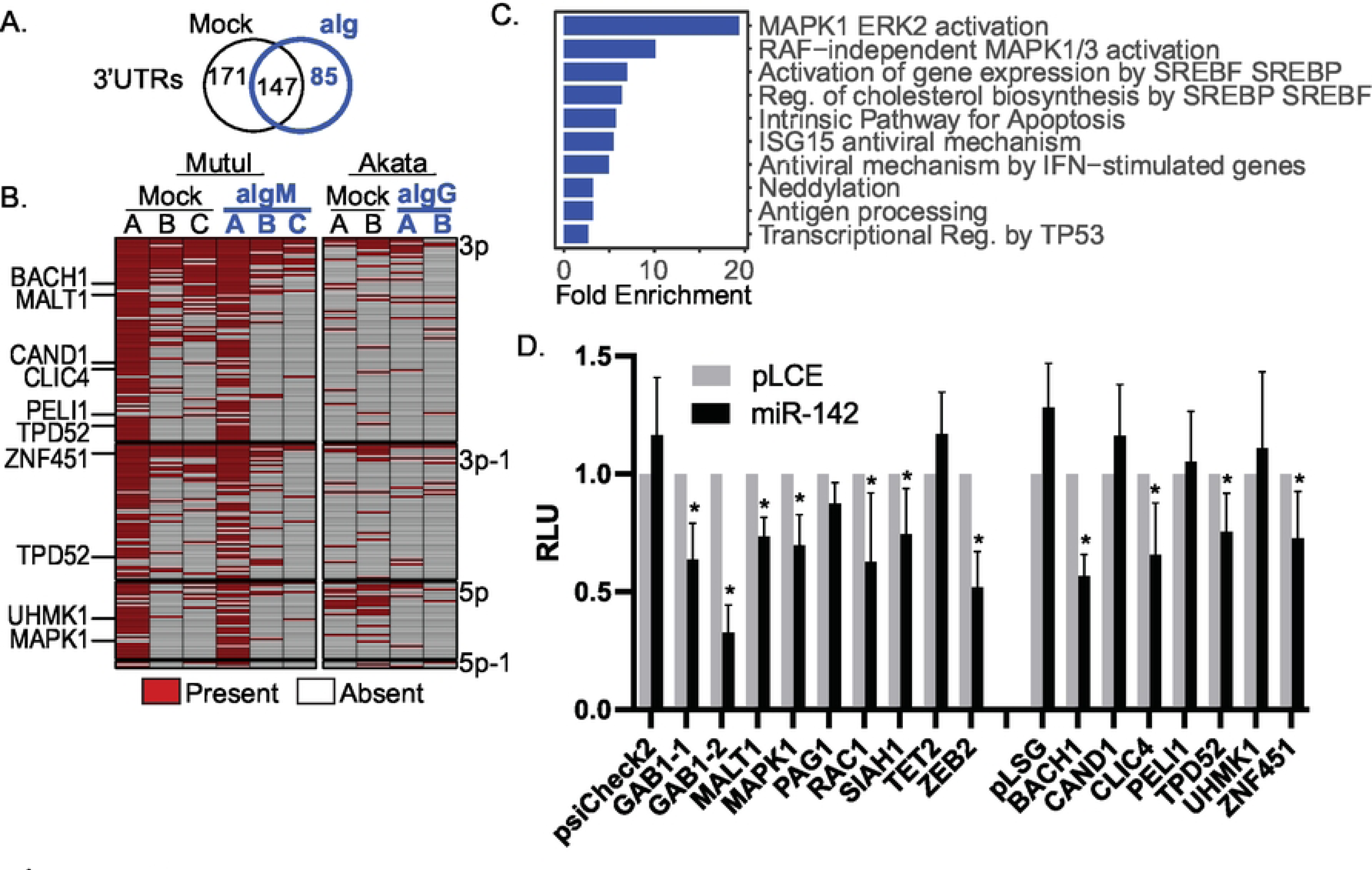
PAR-CLIP analysis identifies miR-142 targets. A. Overlap of miR-142 target 3’UTRs in mock and anti-Ig treated MutuI and Akata cells. 89 3’UTRs targeted by miR-142 miRNAs were identified in reactivated cells. B. miR-142 interactions identified in PAR-CLIP datasets from MutuI and Akata BL cells. Shown are 3’UTR sites that were captured in at least two or more datasets and harbor canonical seed matches (i.e. >=7mer1A) to either miR-142-3p, -3p-1, -5p, or -5p-1 as indicated. 3’UTRs for select genes tested in luciferase assays (panel D.) are highlighted. C. Top ten pathways enriched for miR-142 targets as determined by ShinyGo v0.77 and the Reactome database. D. 293T cells were co-transfected with psiCheck2 vector harboring selected 3’UTR luciferase reporters and miR-142 expression vector (pLCE-based). 72 hrs post-transfection, cells were lysed and assayed for dual luciferase activity. Shown are the averages of at least four experiments performed in triplicate. Student’s t-test, *p<0.05. RLU=relative light units.

miR-142 has various described roles in hematopoiesis, B cell maturation, IL-15 signaling, fatty acid oxidation, and TGF-beta signaling (37, 45, 46, 47); however, its role in EBV infection is not well characterized. To gain an understanding of the major pathways impacted by miR-142 in EBV- infected BL cells, we analyzed the full list of 409 PAR-CLIP-determined miR-142 targets for their enrichment in specific cell signaling pathways using ShinyGO 0.77 and the curated Reactome database (48). The top 10 pathways regulated by miR-142 miRNAs are shown in Figure 3C and included MAPK1/ERK2 signaling, regulation of cholesterol biosynthesis, apoptosis, and p53 transcriptional regulation. We also used Ingenuity pathway analysis (IPA) to interrogate the list of miR-142 targets. Congruent with the Reactome list, top IPA pathways were related to Erk/MAPK signaling, apoptosis, and p53 signaling (Fig. S2A). We then compared our CLIP data to other published Ago-CLIP-seq studies that were performed on B cell lymphoma lines. Of the 409 3’UTR targets identified in our PAR-CLIP analysis for all miR-142 isoforms, 263 overlapped with other CLIP studies (Fig. S2B), while 146 were novel. 56 3’UTR targets were specifically novel for miR- 142-3p (Fig. S2C). Combined assessment of PAR-CLIP and published CLIP data determined that miR-142 targets were enriched in “Diseases of signal transduction by growth factor receptors and second messengers” (Reactome R-HSA-5663202). These observations indicate that, of the multiple genes post-transcriptionally targeted by miR-142 in EBV-infected BL, many are commonly linked to pathways involved in cell growth activated through receptor tyrosine kinases.

To gain further evidence that miR-142 indeed regulates expression of the genes identified by PAR-CLIP, we selected several targets for further investigation based upon their relevance to MAPK signal transduction, including GAB1, MAPK1, MALT1, and the previously documented miR-142 target RAC1 (Rac Family Small GTPase 1) (49). Additional targets were selected for validation based on the fact that they have previously been reported EBV miRNA targets (i.e. BACH1, CLIC4, TPD52, ZNF451) and thus, are likely relevant to EBV latency and/or reactivation (Fig. S2B). 3’UTRs containing canonical miR-142 seed-match sites (i.e. >7mer1A) were cloned into luciferase reporter vectors and tested for activity in the presence of ectopic miR-142. Through this analysis, we found 10 out of 15 3’UTRs that responded to miR-142, confirming these as direct targets (Fig. 3D).

### miR-142 impacts Mek/Erk signaling

Signaling through the BCR activates multiple arms of the MAPK signaling cascade including p38 kinase, JNK (Jun N-terminal kinase), and Erk (extracellular regulated kinase). Erk/MAPK signaling can be activated in response to growth factors or external antigenic triggers and in B cells, these signaling events facilitate key differentiation and maturation processes—such as plasma cell differentiation (50) which initiates EBV lytic replication (8). Given that the MAPK1 (Erk2) 3’UTR reporter was downregulated by miR-142 (Fig. 3D) and that two pathway analysis tools identified miR-142-targeted components related to Erk/MAPK signal transduction (Fig. 3C and S2), we hypothesized that miR-142 might suppress signaling through Erk1/2. A series of phosphorylation events involving Ras/Raf/Mek activate Erk1/2 and Elk-1 (E-twenty six-like transcription factor) downstream of the BCR and other cell surface receptors (51). Erk-mediated phosphorylation of Elk-1 induces its association with the serum response factor (SRF) and subsequent binding to serum response element (SRE) sequences, such as those found within the early growth response 1 (EGR1) promoter (52).

To determine whether miR-142 could regulate the Ras/Raf/Mek/Erk signaling arm, we initially tested a MAPK reporter harboring SRE-responsive luciferase in 293T cells treated with or without epidermal growth factor (EGF) to activate the pathway. Transfection of a miR-142 expression vector (pLCE-miR-142) into these cells significantly reduced SRE activation and luciferase expression in response to EGF (Fig. S3A). We next measured EGR1 transcripts in the two miR- 142 mutant MutuI cell lines that were treated with anti-IgM. Even at 24 and 48 hrs post-treatment, EGR1 levels were increased in miR-142 mutant MutuI cells (Fig. S3C). To investigate in more detail, we subsequently examined protein abundance and phosphorylation status of Mek and Erk at early times post BCR stimulation. While total Mek and Erk levels were relatively unchanged upon disruption of miR-142 (Fig. 4A, C, and E), phosphorylated Mek and to some degree phosphorylated Erk were elevated in miR-142 mutant MutuI cells following stimulation with anti-IgM (Fig. 4A, B, and D). These results demonstrate that miR-142 effectively modulates Mek/Erk signaling in EBV-infected BL cells and moreover, imply that miR-142 functionally regulates expression of components upstream of Mek phosphorylation.

**Figure 4.**
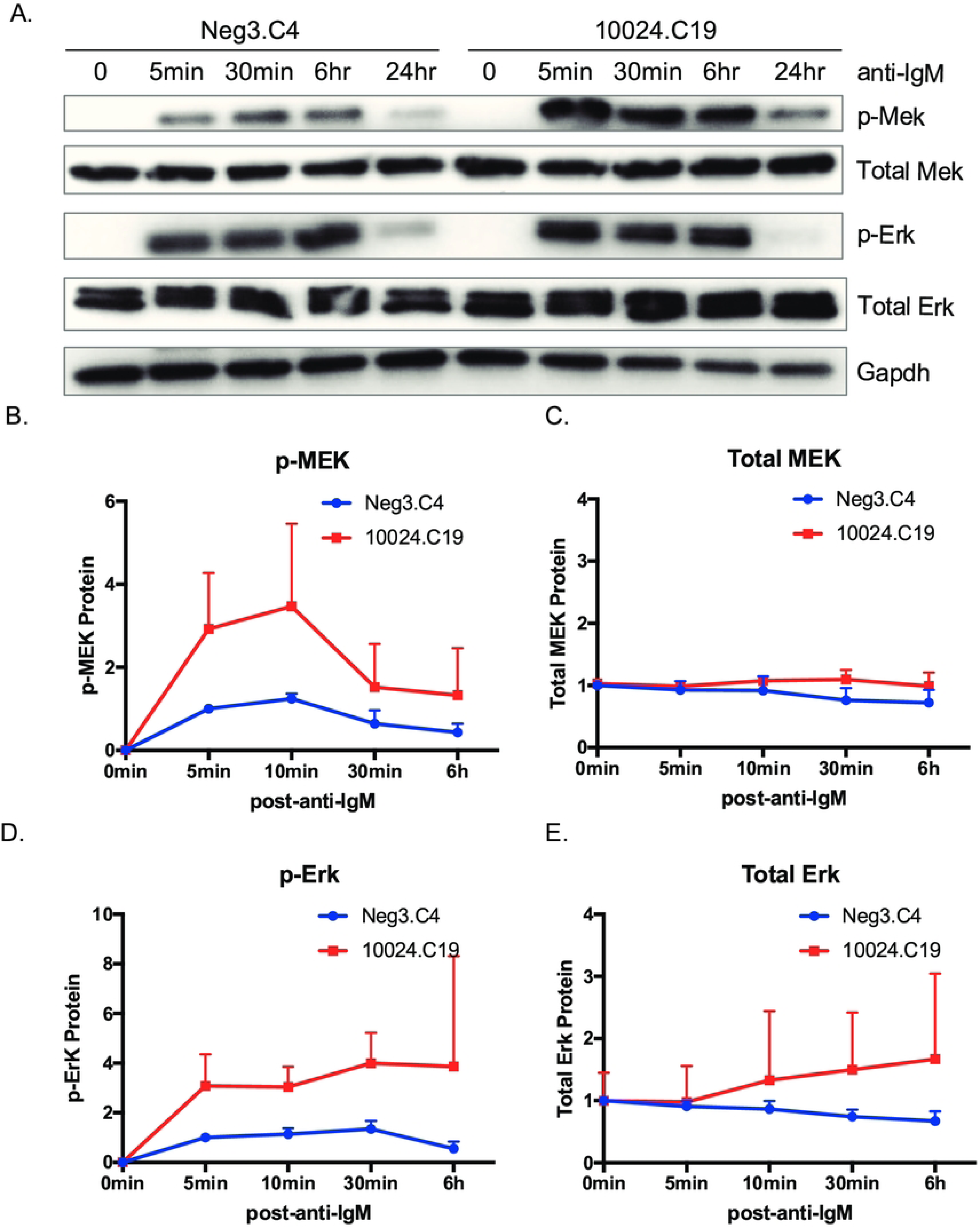
miR-142 regulates Mek/Erk pathway activity in EBV-infected BL cells. A. Time course of Mek/Erk pathway proteins by immunoblot in control (Neg3.C4) or miR-142 mutant (10024.C19) MutuI cells following anti-IgM treatment. Cells were serum starved for 2 hrs prior to anti-IgM treatment for the indicated times. Gapdh levels are shown as loading controls. Shown is the representative of three independent experiments. B through E. Band intensities in panels A and B were quantified using ImageJ and normalized to loading controls. Phospho-proteins were reported relative to 5 min anti-IgM treated Neg3.C4 cells, and total proteins were reported relative to Neg3.C4 cells at 0 min.

### Small molecule inhibitors of either Mek or Raf partially rescue lytic phenotypes observed in miR-142 mutant cells

To further investigate the Mek/Erk signaling pathway in miR-142 mutant cells, we tested small molecule inhibitors to Mek1/2 (cobimetinib) or to Raf (dabrafenib) which is directly upstream of Mek (Fig. 5A). Cobimetinib (GDC-0973) is a selective inhibitor of Mek kinase and blocks Mek-mediated signaling (53). Dabrafenib is a BRAF inhibitor that is FDA-approved for treatment of solid tumors (54). miR-142 mutant MutuI cells were pre-treated with either inhibitor and subsequently induced to reactivate. Phosphorylated Erk and phosphorylated Mek were substantially reduced (Fig. 5B and C) upon treatment, confirming activity of the inhibitors. Assessment of EBV lytic proteins revealed lower levels of Zta in the presence of either cobimetinib or dabrafenib compared to mock treated cells (Fig. 5D and E). These results demonstrate that specific inhibitors to either Mek or Raf can partly attenuate EBV reactivation responses in miR- 142 mutant cells, but do not completely block the lytic cycle.

**Figure 5.**
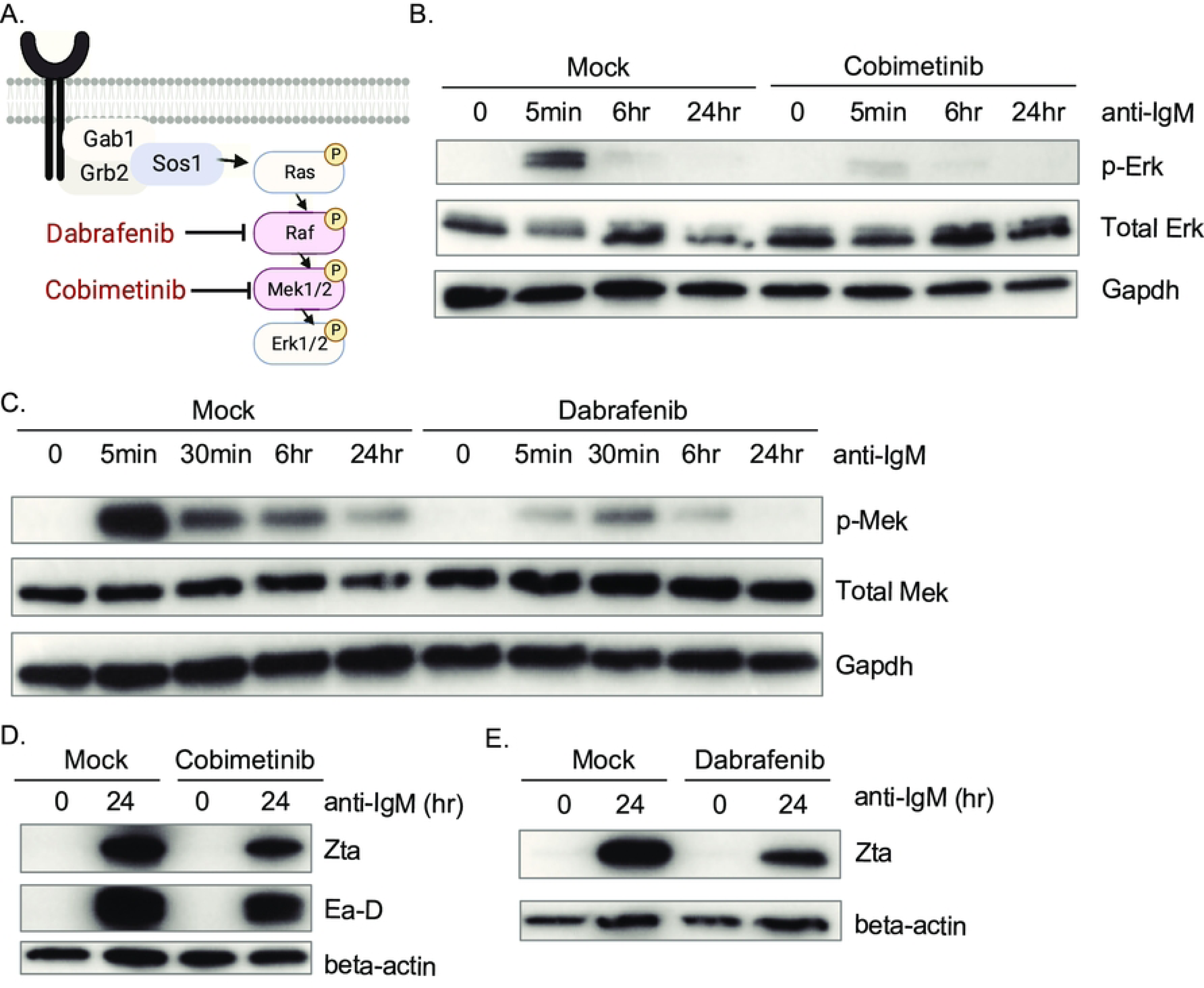
Chemical inhibition of the Raf/Mek/Erk pathway limits lytic reactivation in miR- 142 mutant cells. A. Schematic illustration of the Ras/Raf/Mek/Erk kinase cascade and the protein targets of dabrafenib and cobimetinib. B. Immunoblot validation of cobimetinib activity in miR-142 mutant MutuI cells (10024.C19) in blocking phosphorylation of Erk by Mek. Gapdh levels are shown as loading controls. C. Immunoblot validation of dabrafenib activity in miR-142 mutant MutuI cells (10024.C19) in blocking phosphorylation of Mek by Raf. Gapdh levels are shown as loading controls. D. and E. EBV lytic proteins are reduced in miR-142 mutant MutuI cells upon inhibition of the Raf/Mek/Erk pathway. 10024.C19 cells were treated with either cobimetinib or dabrafenib followed by anti-IgM for 24 hrs. Zta and Ea-D were assayed by immunoblot. Beta-actin levels are shown as loading controls. Shown are representative images of at least three independent experiments.

### Ras guanine exchange factor SOS1 is targeted by miR-142

Given that Raf and Mek inhibitors partially rescued phenotypes associated with miR-142 perturbation, we hypothesized that the host factors upstream of Raf are most likely contributing to the enhanced EBV reactivation observed upon loss of miR-142. Ras-GTPases activate the serine/threonine kinase Raf, and upstream of Ras-GTP are Ras guanine exchange factors such as SOS1 (son of sevenless 1) (55). Notably, the SOS1 3’UTR contains at least six predicted binding sites for miR-142-3p and miR-142-5p (Fig. 6A) and has previously been reported as a miR-142 target (41). We therefore hypothesized that loss of miRNA-mediated SOS1 suppression might augment early activation steps in the BCR signaling cascade. To confirm that the SOS1 3’UTR responds to miR-142, we tested a luciferase reporter against either pLCE-miR-142 or pLCE-miR-142-3p (Fig. 6B, Fig. S3B). Luciferase activity was reduced in both conditions, showing that miR-142 directly impacts the SOS1 3’UTR. We next measured SOS1 protein and RNA levels in miR-142 mutant cells. RAC1, a previously validate miR-142 target, and SIAH1 transcripts were tested in parallel (Fig. 6C). Pertubation of miR-142 in MutuI cells resulted in increased SOS1 expression at both the RNA and protein levels (Fig. 6C-E), confirming that miR-142 regulates SOS1 in EBV-infected BL cells.

**Figure 6.**
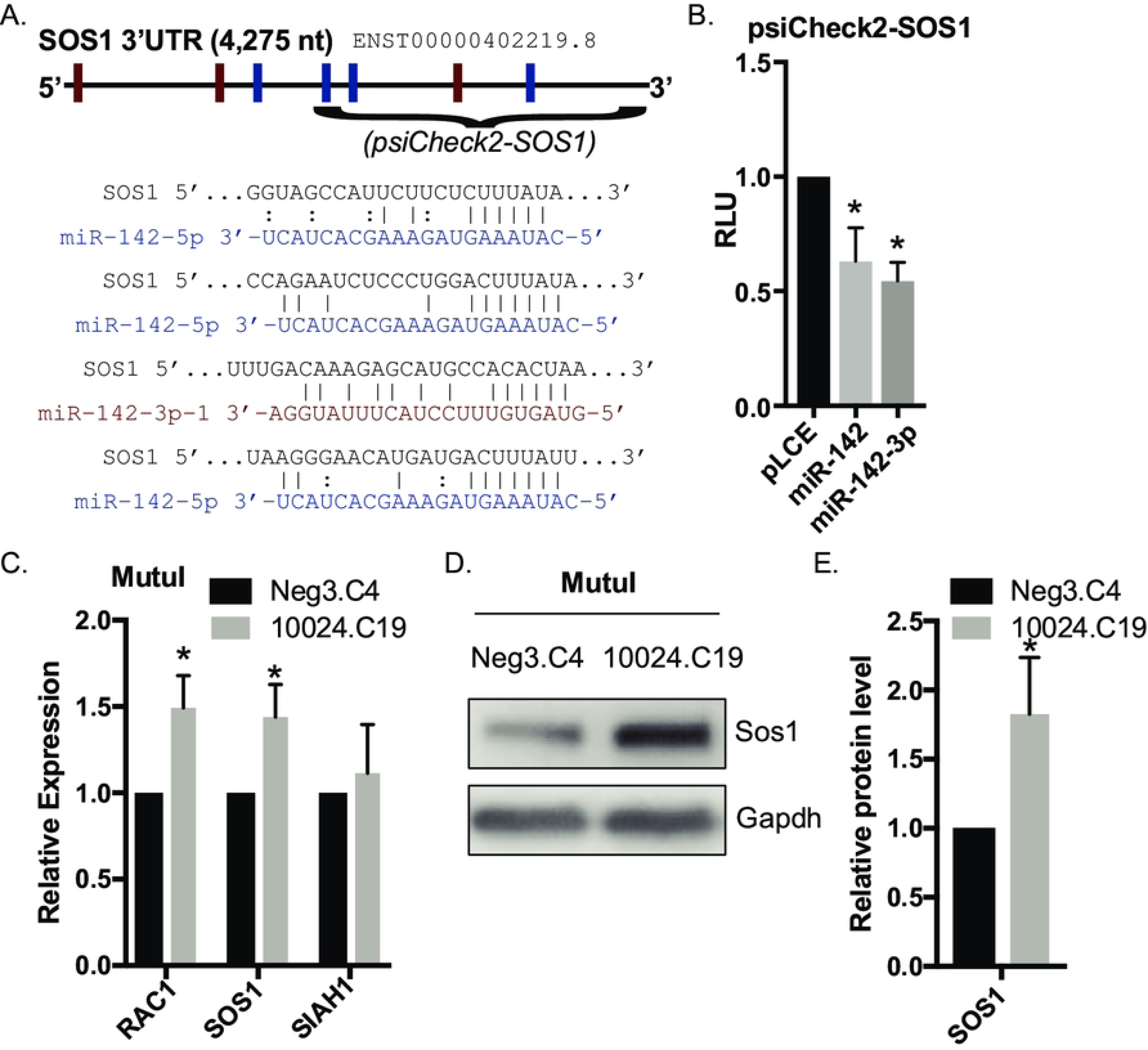
SOS1 is targeted by miR-142. A. Schematic of the SOS1 3’UTR with miR-142-3p and miR-142-5p binding sites marked with red and blue rectangles, respectively. The portion of the SOS1 3’UTR that is cloned into psiCheck2 and the four predicted miR-142 binding sites located within this region are highlighted. B. 293T cells were co-transfected with the psiCheck2-SOS1 luciferase reporter and either pLCE control or pLCE-based expression vectors for miR-142-3p or all miR-142 miRNAs. 72 hrs post-transfection, cells were lysed and assayed for dual luciferase activity. Shown are the averages of six experiments performed in triplicate. By Student’s t-test, *p<0.05. RLU=relative light units. C. RAC1 and SOS1 mRNA levels increase in the absence of miR-142. Total RNA was collected from miR-142 mutant MutuI cells (10024.C19) and control cells (Neg3.C4). RAC1, SOS1 and SIAH1 expression was determined by qRT-PCR. Values are normalized to GAPDH and shown relative to control cells. Shown is the average of three independent experiments. By Student’s t- test, *p<0.05. D. and E. Immunoblot analysis and quantification of Sos1 protein in control (Neg3.C4) and miR-142 mutant (10024.C19) MutuI cells. Gapdh levels are shown as loading controls. Band intensities were quantified using ImageJ, normalized to loading controls, and reported relative control cells. Shown is the representative of four independent experiments. By Student’s t-test, *p<0.05.

### Inhibition of SOS1 Limits EBV Lytic Reactivation

Since SOS1 protein expression was upregulated in the absence of miR-142 and correlated with increased lytic EBV gene products, we investigated whether knocking down SOS1 would limit lytic reactivation in response to anti-IgM. To phenocopy RNAi-mediated inhibition, shRNAs against SOS1 were stably introduced into miR-142 mutant MutuI cells. Following anti-IgM stimulation, we found lower levels of lytic proteins (Fig. 7A) as well as reduced phosphorylation of Mek and Erk (Fig. 7B) in cells lacking SOS1 compared to pLCE controls. Similar effects were observed when cells were pre-treated with a small molecule inhibitor to SOS1 (BI-3406) and then induced into the lytic cycle (Fig. 7C and D). Taken together, these results show that SOS1 activity is needed for EBV reactivation by BCR crosslinking and moreoever, that miR-142 restricts lytic reactivation in part by targeting SOS1.

**Figure 7.**
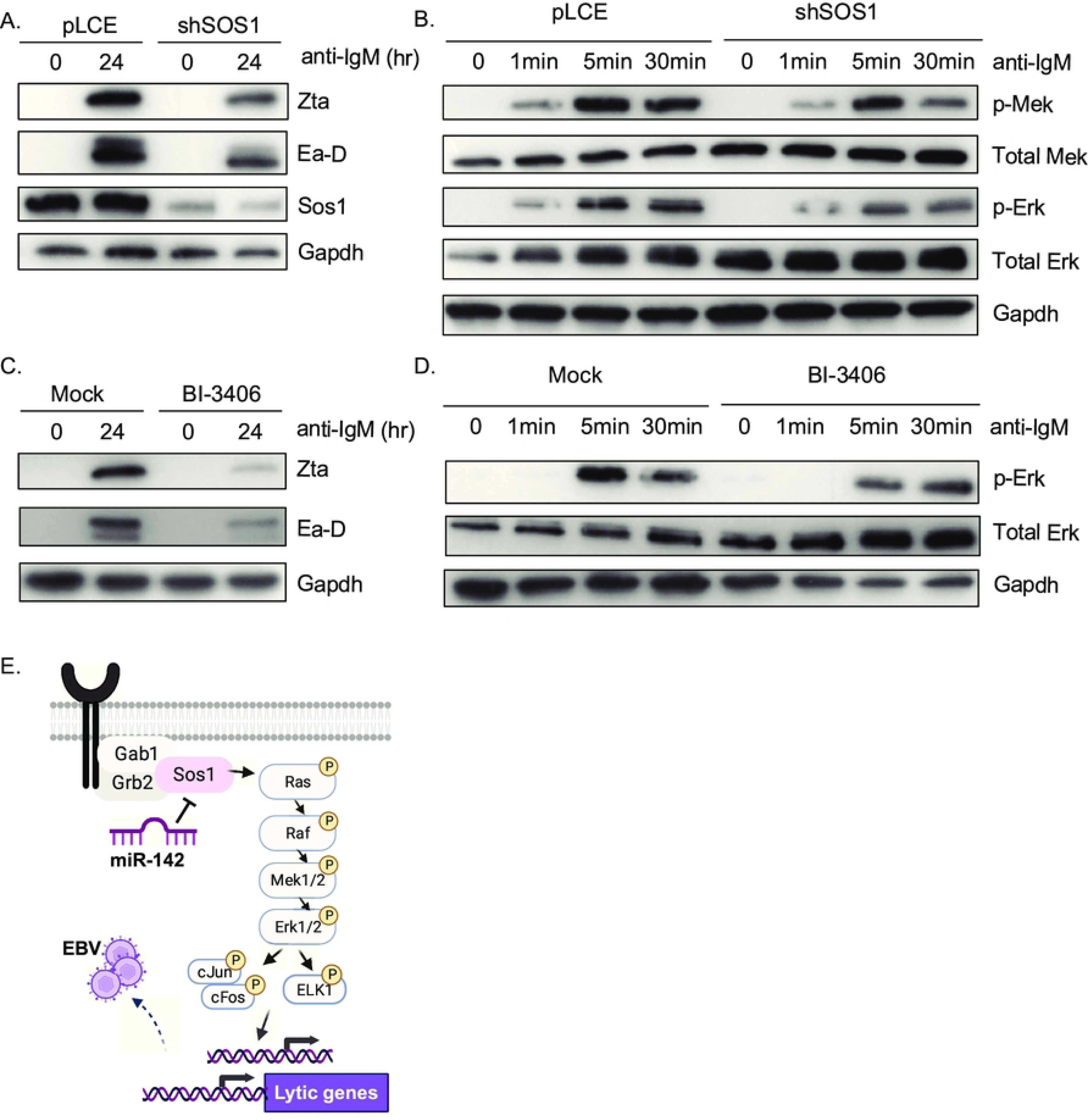
Abrogation of SOS1 limits the enhanced lytic reactivation by miR-142 knockdown. A. RNAi-mediated inhibition of SOS1 attenuates EBV lytic gene expression in response to BCR crosslinking. miR-142 mutant MutuI 10024.C19 cells were stably transduced with empty vector control (pLCE) or shRNA against SOS1 and treated with anti-IgM for 0 or 24 hrs. Immunoblots were performed on lysates for lytic gene products and for Sos1 as knockdown validation. Gapdh levels are shown as loading controls. Shown is the representative of three independent experiments. B. Inhibition of SOS1 reduces activation of the Ras/Raf/Mek/Erk pathway. Time course of Mek/Erk pathway proteins by immunoblot in miR-142 mutant MutuI cells transduced with empty vector (pLCE) or shSOS1 following anti-IgM treatment. Gapdh levels are shown as loading controls. C. Chemical inhibition of Sos1 attenuates EBV lytic gene expression. EBV lytic proteins were tested by immunoblot in miR-142 mutant MutuI cells pre-treated the Sos1 inhibitor BI-3406 for 24 hrs, followed by anti-IgM treatment for 24 hrs. Gapdh levels are shown as loading controls. E. Erk phosphorylation is reduced in the presence of the Sos1 chemical inhibitor. Time course of Erk proteins tested by immunoblot in miR-142 mutant MutuI cells pre-treated with the Sos1 inhibitor BI-3406 for 24 hrs, followed by anti-IgM treatment for indicated times. Gapdh levels are shown as loading controls. F. Model of how miR-142 regulates induction of EBV lytic gene expression through the Sos1/Ras/Raf/Mek/Erk signaling cascade.

## Discussion

The EBV lytic phase is essential for viral transmission to naïve cells and new hosts. While several studies have investigated roles for viral miRNAs in EBV latency stages and reactivation, there is less knowledge about the activities of host miRNAs during the EBV latent to lytic switch. In this study, we aimed to determine whether specific host miRNAs could regulate EBV reactivation and/or progression of the lytic phase. Through a genome-wide CRISPR/Cas9 miRNA screen, we found that perturbing expression of the majority of host miRNAs had no measurable impact on EBV reactivation (Fig. 1). Genetic ablation of miR-142 in EBV-positive BL cells, however, led to significantly heightened lytic gene expression and enhanced activity of Erk/MAPK signaling in response to BCR stimulation (Fig. 2). Functional dissection of molecular targets revealed that miR-142 directly regulates the SOS1/Ras/Raf/Mek/Erk signaling axis that can be induced by anti- immunoglobulins (Fig. 7). These observations lead us to conclude that miR-142 functionally limits initial triggering of the latent to lytic switch by inhibiting factors driving the MAPK pathway and thereby acts as a restriction factor for EBV reactivation.

The multiple miRNAs expressed from the miR-142 locus have complex relationships with g- herpesviruses. For example, HSURs 1, 2, and 5, which are non-coding RNAs expressed by a g- herpesvirus that infects New World monkeys (herpesvirus saimiri (HVS)), harbor sequence-specific binding sites for miR-142-3p within their 5’ ends (56). Base-pairing between HSUR2 and miR-142-3p enhances HSUR2-mediated repression of target mRNAs such as RB1 and results in suppression of host factors involved in apoptosis (56). In previous studies, the EBV BHRF1 transcript was shown to harbor a binding site for miR-142-3p and miRNA mimics could suppress ectopic BHRF1 protein expression in HEK293T cells (57, 58), suggesting a canonical miRNA interaction with the BHRF1 3’UTR. The functional significance of this interaction in the context of EBV infection has not been fully investigated. In our experiments, we did observe significantly increased levels of BHRF1 protein in miR-142 mutant cells compared to controls (Fig. 2); however, these changes were also accompanied by increased EBV Zta and Ea-D proteins, indicating observed effects are linked to augmented lytic replication and not necessarily due to direct miRNA interactions with the BHRF1 3’UTR or other transcripts arising from the BHRF1 locus. In addition to usurping cellular miR-142 for g-herpesvirus-mediated activities, viruses such as Kaposi’s sarcoma-associated herpesvirus (KSHV) and several non-human primate (NHP) rhadinoviruses encode viral miRNA mimics of miR-142-3p (41, 59, 60). KSHV miR-K10a+1, which exhibits an identical seed sequence to the miR-142-3p-1 5’ isomiR, can regulate a subset of miR-142 targets, enabling the virus to hijack evolutionarily conserved miR-142 regulatory networks (41).

Beyond g-herpesviruses, miR-142-3p has been shown to uniquely restrict replication of eastern equine encephalitis virus (EEEV) in cells of the myeloid lineage by directly binding sites within the 3’UTR of the EEEV RNA viral genome (61). Studies in mouse models with targeted deletion of miR-142 resulted in expansion of CD19+ B cells and elevated BAFF-R (B-cell–activating factor receptor), indicating a role in B cell development and humoral responses (37). miR-142-deficient mice further exhibit defective IL-15 signaling and natural killer cell loss (46) as well as compromised responses to virus infection, including increased susceptibility to herpes simplex virus (HSV-1) or murine cytomegalovirus (mCMV) infections (37, 46). Thus, adequate miR-142 function is necessary for maintaining intact immune cell responses and controlling replication of many different viruses.

Mutations in the human miR-142 locus that disrupt expression of miR-142-3p and miR-142-5p have been frequently observed in cancers. Assessment of somatic mutations in miRNA loci in TCGA samples revealed that the highest frequency of mutations in DLBCLs occurred in the miR- 142 locus (35). In published miRNA profiling studies, miR-142-5p was found to be upregulated in EBV-negative BL cases compared to EBV positive cases (39), prompting us to measure miRNA levels in various transformed B cell lines, including DLBCLs, LCLs, and BL (Fig. 1). Surprisingly, we found no remarkable differences in miR-142 expression in EBV-positive versus EBV-negative BL cell lines; however, both miR-142-5p and miR-142-3p were detected at higher levels in DLBCLs and BLs compared to latency III LCLs. Such findings support the idea that cell context— such as B cell maturation state—contributes to miR-142 expression.

Through PAR-CLIP analysis to investigate the miRNA targetome, we identified >400 Ago-bound 3’UTR targets of miR-142 in EBV-positive BL cells (Fig. 3). Interestingly, ∼40% of miR-142 interactions were captured only in mock treated cells while ∼20% of interactions were found only in anti-Ig treated cells, highlighting the dynamic nature of miRNA interactions influenced by cellular responses to external stimuli. Validated interactions for miR-142 included targets captured only in mock treated cells (MALT1, SIAH1, ZEB2), only in anti-Ig treated cells (PAG1), or in both conditions (MAPK1, BACH1, CLIC4, TPD52, ZNF451). Further investigation is warranted to understand the biologically important interactions that are captured under specific environmental conditions, such as in response to BCR engagement.

In summary, our findings shed new mechanistic insight into the functional role of host miR-142 in orchestrating the EBV latent to lytic switch. Significantly, through these studies, we found that small molecule inhibitors to either SOS1, Raf, or Mek can dampen EBV lytic reactivation. Moreover, our experiments reveal that, despite the loss of miR-142 through CRISPR engineering, functional effects may be restored by pharmacological manipulation of miR-142 targeted pathways. These findings have important implications for cancers in which miR-142 is affected.

## Methods

### Cell culture, lentivirus, and transductions

B cell lines were maintained at 37°C in a 5% CO2-humidified atmosphere in RPMI 1640 supplemented with 10% fetal bovine serum (FBS) and 1% penicillin, streptomycin, and L-glutamine (P/S/G). HEK293T cells were maintained in high-glucose Dulbecco modified Eagle medium supplemented with 10% FBS and 1% P/S/G. For preparation of lentiviruses, HEK293T cells were plated in 15-cm plates in complete media and transfected using polyethylenimine (PEI) with 15 μg of lentivector, 9 μg of pDeltaR8.75, and 6 μg of pMD2G. Medium was changed to complete RPMI 1640 at between 8 and 16 h post-transfection. Lentiviral particles were harvested by sterile filtration of the supernatant using a 0.22-μm filter at 48 and 96 h post-transfection and used to transduce ca. 1x10^6^ to 5x10^6^ cells. For BCR cross-linking, BL cells were spun down and plated at 0.5x10^6^ cells in fresh medium containing soluble anti-IgM (Sigma) at the concentration of 5 μg/ml and analyzed at times indicated.

### Plasmids

Inducible Cas9 vector pCW-Cas9-blast and gRNA expression vector pLentiGuide-puro were purchased from Addgene. Oligonucleotide sequences used for gRNA cloning are available upon request. pLCE-miR-142 and pLCE-miR-142-3p vectors were generously provided by Dr. Eva Gottwein and Dr. Mark Manzano from Northwestern University (41). miRNA expression was confirmed by Taqman qRT-PCR (Fig. S3B). psiCheck2 and pLSG 3′ UTR constructs are previously described (26).

### Human miRNA CRISPR/Cas9 screen and data analysis

The gRNA library targeting >1500 human miRNA loci was obtained from Addgene (32). MutuI cells were transduced with lentiparticles derived from the doxycycline inducible Cas9 vector pCW-Cas9 (Addgene Plasmid #50661) and selected for stable integration with 2 μg/mL puromycin. Resulting MutuI-pCW-Cas9 cells were then transduced with the pLX-miR library (32), a gift from Dr. Ren-Jang Lin (Addgene Pooled Library #112200) and selected for stable integration with 5 μg/mL blasticidin. To verify library composition, resulting MutuI-pCW-Cas9-LX-miR cells were used to produce gRNA targeted sequencing libraries and sequenced on an Illumina MiSeq system. The original pLX-miR plasmid pool was deep sequenced in parallel. For screening purposes, three replicates of 2x10^7^ MutuI-pCW-Cas9-LX-miR cells were resuspended at 10^6^ cells per mL in normal growth media supplemented with 10 μg/mL anti-IgM. Forty-eight hours later, cells were collected by centrifugation and resuspended in 4 mL ice cold phosphate-buffered saline (PBS). 16 mL of ice-cold methanol was added dropwise while vortexing to fix cells and then incubated on ice for 20 minutes. Cells were washed in 1 mL Cell Sort Media (PBS, 0.05M EDTA + 2% BSA) and collected by centrifugation at 500xg for 3 minutes. Cells were resuspended in 1 mL Cell Staining Buffer. 100 μL Human TruStain FcX™ (BioLegend) was added, gently mixed, and the cells were incubated on ice for 10 minutes. 20 μL of anti-gp350 antibody was added and gently mixed. Tubes were incubated on ice for 30 minutes. Cells were then washed three times with 1 mL Cell Staining Buffer, resuspended in 1 mL Cell Staining Buffer with 5 μL of secondary Alexa Fluor 488 antibody, and incubated for 30 minutes on ice in the dark. Cells were subsequently washed three times with 1 mL Cell Staining Buffer and finally resuspended in 2 mL Cell Sort Media. Cells were sorted into gp350 positive and negative populations using a BD FACS Aria-II cell sorter, pelleted, and DNA was extracted using the Qiagen Blood and Tissue kit according to the manufacturer’s protocol. gRNA target sequences were amplified using a two- step approach adapted from the TKOv3 CRISPR library protocol (62). gRNA regions were first amplified from genomic DNA using published lentiC-dseq-2-F1 and LX-dseq-R1 primers (32). A second round of PCR was performed on these amplicons using a staggered pooled primer approach to add Illumina adaptor and index sequences for sequencing. Libraries were size fractionated on agarose gels and QC was performed with an Agilent BioAnalyzer. Resulting libraries were sequenced on an Illumina NovaSeq at the OHSU MPSSR.

Following sequencing, adaptor sequences were trimmed from reads using cutadapt (63). Reads were then mapped to the pLX-miR library reference sequences using Bowtie2 (64). Individual guide counts per miRNA were summarized with SAMtools (65). Replicates were combined; subsequent normalization, fold change calculations, and statistical testing was performed with the drugZ software (33). To reduce non-miRNA-mediated off target hits, analysis was limited to the cellular miRNAs that were previously identified in MutuI cells (26).

### Generation of individual miR-142 mutant cell lines by CRISPR/Cas9

Two separate guide sequences targeting the miR-142 locus (SgID #10024 and #10033) from the pLX-miR pool (32) were cloned into vector pBA904 (Addgene Plasmid #122238). MutuI cells were transduced with lentiparticles derived from pCW-Cas9-Blast (Addgene Plasmid #83481) and selected for stable integration with 10 μg/mL blasticidin. Resulting MutuI-Cas9-Blast cells were then transduced with lentiparticles derived from pBA904-10024 or pBA904-10033 and selected for stable integration with 1 μg/mL puromycin. Individual clones were derived by plating cells by limiting dilution into 96-well plates. Resulting clones were expanded and subsequently screened by qRT-PCR to identify cell lines in which miR-142 expression was disrupted.

### PAR-CLIP and bioinformatics analysis

MutuI and Akata cells were expanded in log phase, then cultured in 100 uM 4-SU (4-thiouridine) for 2 hrs prior to addition of 5 μg/mL anti-IgM or anti-IgG. Cells were subsequently cultured for 20 hrs, crosslinked at UV 365 nm, and cell pellets stored at –80*C. To immunopurify Ago-bound RNAs, cell pellets were lysed in NP40 lysis buffer and RNAs were immunopurified using antibodies to Ago2 (Abcam) as previously described (42). cDNA libraries were sequenced on the Illumina platform. Raw sequencing reads were obtained in FASTQ format. FASTQ files were concatenated, adapters removed, and sequences collapsed using the FASTX-toolkit (http://hannonlab.cshl.edu/fastx_toolkit/), removing reads that contained an N or were <12 nt in length after adapter removal. Processed reads were aligned to the human genome (HG38) with Bowtie (66). Mapped reads were then run through PARalyzer (43), requiring at least three reads per cluster. The PARalyzer distribution file was subsequently run through microMUMMIE (44) to predict additional Ago interaction sites. Ago interaction sites aligning to 3’UTRs of human protein-coding transcripts were annotated for canonical miR-142 seed matches (>=7mer1A).

### Luciferase assays

Dual luciferase reporter assays were performed as previously described (26). Briefly, HEK293T cells were plated in 96-well black-well plates and co-transfected with 20 ng of 3′ UTR reporter and 250 ng of control vector (pLCE) or miRNA expression vector using Lipofectamine 2000 (Thermo Fisher). Cells were harvested in 1x passive lysis buffer (Promega) at 48-72 hrs post-transfection and lysates were assayed for luciferase activity using the dual-luciferase reporter assay system (Promega) and a luminometer. All values are reported as relative light units (RLU) relative to luciferase internal control and normalized to pLCE control vector.

### RNA isolation and qRT-PCR

Total RNA was extracted using TRIzol (Thermo Fisher) according to the manufacturer’s protocol, except substituting the wash step with 95% ice cold ethanol. RNA concentration and integrity were analyzed using a NanoDrop. To assess gene expression, RNA was DNase-treated and reversed transcribed using MultiScribe (Thermo Fisher) with random hexamers. Cellular and viral genes were detected using qPCR primers to gene-specific regions and PowerUp SYBR green qPCR mastermix (Thermo Fisher). All genes were normalized to GAPDH levels. To assay miRNA expression, total RNA was reverse transcribed using miRNA-specific Taqman (Thermo Fisher) primers and miRNAs were detected using miRNA-specific Taqman probes and Taqman Universal PCR mastermix (Thermo Fisher). All miRNAs were normalized to hsa-miR-16. Oligonucleotides sequences are available upon request. Viral loads were determined by qPCR for LMP1 copies using a standard curve. All PCR reactions were performed in technical replicates (duplicates or triplicates).

### Immunoblotting

Cells were lysed in RIPA lysis buffer. Protein concentrations were determined using the bicinchoninic acid protein assay kit (Thermo Scientific), and 10-20μg of total protein lysate was resolved on 5-20% Tris-glycine SDS-PAGE and then transferred onto Immobilon polyvinylidene difluoride membranes. Blots were probed with primary antibodies to SOS1 (5890S; Cell Signaling), phospho-Mek (9154S; Cell Signaling), Mek (9126S; Cell Signaling), phospho-Erk (4370S; Cell Signaling), Erk (4695S; Cell Signaling), EBV EaD (BMRF1, sc-58121; Santa Cruz), EBV Zebra (BZLF1, sc-53904; Santa Cruz), BHRF1 (biorbyt orb518144), GAPDH (sc-47724; Santa Cruz), or beta-actin (sc-47778; Santa Cruz), followed by horseradish peroxidase-conjugated secondary antibodies (anti-rabbit IgG or anti-mouse IgG). Blots were developed with enhanced chemiluminescent substrate (Pierce). Band intensities were quantified using ImageJ, normalized to loading controls, and reported relative to control cells.

### Datasets

Sequencing datasets are available through NCBI Sequence Read Archive (SRA) under BioProject PRJNA796351 and PRJNA1030428.

### Statistical analysis

Statistical analysis was performed using GraphPad Prism 9.0.1 or Microsoft Excel. For luciferase and qRT-PCR assays, Student’s t-tests were used to determine statistical significance and p-values less than 0.05 were considered significant.

## Acknowledgements

This study was supported by grant R01 AI143620 from the National Institute of Allergy and Infectious Diseases to RLS.

## References

1. Thorley-Lawson DA. EBV Persistence--Introducing the Virus. Curr Top Microbiol Immunol. 2015;390(Pt 1):151–209.

2. Balfour HH, Jr., Odumade OA, Schmeling DO, Mullan BD, Ed JA, Knight JA, et al. Behavioral, virologic, and immunologic factors associated with acquisition and severity of primary Epstein-Barr virus infection in university students. J Infect Dis. 2013;207(1):80-8.

3. Farrell PJ. Epstein-Barr Virus and Cancer. Annu Rev Pathol. 2019;14:29–53.

4. Bjornevik K, Cortese M, Healy BC, Kuhle J, Mina MJ, Leng Y, et al. Longitudinal analysis reveals high prevalence of Epstein-Barr virus associated with multiple sclerosis. Science. 2022;375(6578):296-301.

5. Moormann AM, Bailey JA. Malaria - how this parasitic infection aids and abets EBV- associated Burkitt lymphomagenesis. Curr Opin Virol. 2016;20:78–84.

6. Khan G, Hashim MJ. Global burden of deaths from Epstein-Barr virus attributable malignancies 1990-2010. Infect Agent Cancer. 2014;9(1):38.

7. Kenney SC, Mertz JE. Regulation of the latent-lytic switch in Epstein-Barr virus. Semin Cancer Biol. 2014;26:60–8.

8. Laichalk LL, Thorley-Lawson DA. Terminal differentiation into plasma cells initiates the replicative cycle of Epstein-Barr virus in vivo. J Virol. 2005;79(2):1296–307.

9. Rosemarie Q, Sugden B. Epstein-Barr Virus: How Its Lytic Phase Contributes to Oncogenesis. Microorganisms. 2020;8(11).

10. Deng Y, Munz C. Roles of Lytic Viral Replication and Co-Infections in the Oncogenesis and Immune Control of the Epstein-Barr Virus. Cancers (Basel). 2021;13(9).

11. Tierney RJ, Shannon-Lowe CD, Fitzsimmons L, Bell AI, Rowe M. Unexpected patterns of Epstein-Barr virus transcription revealed by a high throughput PCR array for absolute quantification of viral mRNA. Virology. 2015;474:117–30.

12. Donati D, Espmark E, Kironde F, Mbidde EK, Kamya M, Lundkvist A, et al. Clearance of circulating Epstein-Barr virus DNA in children with acute malaria after antimalaria treatment. J Infect Dis. 2006;193(7):971–7.

13. Ji MF, Wang DK, Yu YL, Guo YQ, Liang JS, Cheng WM, et al. Sustained elevation of Epstein- Barr virus antibody levels preceding clinical onset of nasopharyngeal carcinoma. Br J Cancer. 2007;96(4):623–30.

14. Stevens SJ, Verkuijlen SA, Hariwiyanto B, Harijadi, Paramita DK, Fachiroh J, et al. Noninvasive diagnosis of nasopharyngeal carcinoma: nasopharyngeal brushings reveal high Epstein-Barr virus DNA load and carcinoma-specific viral BARF1 mRNA. Int J Cancer. 2006;119(3):608–14.

15. Su Y, Yuan D, Chen DG, Ng RH, Wang K, Choi J, et al. Multiple early factors anticipate post- acute COVID-19 sequelae. Cell. 2022;185(5):881–95 e20.

16. Klein J, Wood J, Jaycox J, Lu P, Dhodapkar RM, Gehlhausen JR, et al. Distinguishing features of Long COVID identified through immune profiling. medRxiv. 2022.

17. Peluso MJ, Deveau TM, Munter SE, Ryder D, Buck A, Beck-Engeser G, et al. Chronic viral coinfections differentially affect the likelihood of developing long COVID. J Clin Invest. 2023;133(3).

18. Rahman MA, Kingsley LA, Breinig MK, Ho M, Armstrong JA, Atchison RW, et al. Enhanced antibody responses to Epstein-Barr virus in HIV-infected homosexual men. J Infect Dis. 1989;159(3):472–9.

19. Reusch JA, Nawandar DM, Wright KL, Kenney SC, Mertz JE. Cellular differentiation regulator BLIMP1 induces Epstein-Barr virus lytic reactivation in epithelial and B cells by activating transcription from both the R and Z promoters. J Virol. 2015;89(3):1731–43.

20. Bhende PM, Dickerson SJ, Sun X, Feng WH, Kenney SC. X-box-binding protein 1 activates lytic Epstein-Barr virus gene expression in combination with protein kinase D. J Virol. 2007;81(14):7363–70.

21. Sun CC, Thorley-Lawson DA. Plasma cell-specific transcription factor XBP-1s binds to and transactivates the Epstein-Barr virus BZLF1 promoter. J Virol. 2007;81(24):13566–77.

22. Wen Y, Jing Y, Yang L, Kang D, Jiang P, Li N, et al. The regulators of BCR signaling during B cell activation. Blood Sci. 2019;1(2):119–29.

23. Xu W, Berning P, Lenz G. Targeting B-cell receptor and PI3K signaling in diffuse large B-cell lymphoma. Blood. 2021;138(13):1110–9.

24. Iwakiri D, Takada K. Phosphatidylinositol 3-kinase is a determinant of responsiveness to B cell antigen receptor-mediated Epstein-Barr virus activation. J Immunol. 2004;172(3):1561–6.

25. Chen Y, Fachko D, Ivanov NS, Skinner CM, Skalsky RL. Epstein-Barr virus microRNAs regulate B cell receptor signal transduction and lytic reactivation. PLoS Pathog. 2019;15(1):e1007535.

26. Chen Y, Fachko DN, Ivanov NS, Skalsky RL. B Cell Receptor-Responsive miR-141 Enhances Epstein-Barr Virus Lytic Cycle via FOXO3 Inhibition. mSphere. 2021;6(2).

27. Ellis-Connell AL, Iempridee T, Xu I, Mertz JE. Cellular microRNAs 200b and 429 regulate the Epstein-Barr virus switch between latency and lytic replication. J Virol. 2010;84(19):10329–43.

28. Yin Q, Wang X, Fewell C, Cameron J, Zhu H, Baddoo M, et al. MicroRNA miR-155 inhibits bone morphogenetic protein (BMP) signaling and BMP-mediated Epstein-Barr virus reactivation. J Virol. 2010;84(13):6318–27.

29. Lin Z, Wang X, Fewell C, Cameron J, Yin Q, Flemington EK. Differential expression of the miR-200 family microRNAs in epithelial and B cells and regulation of Epstein-Barr virus reactivation by the miR-200 family member miR-429. J Virol. 2010;84(15):7892–7.

30. Forte E, Salinas RE, Chang C, Zhou T, Linnstaedt SD, Gottwein E, et al. The Epstein-Barr virus (EBV)-induced tumor suppressor microRNA MiR-34a is growth promoting in EBV-infected B cells. J Virol. 2012;86(12):6889–98.

31. Linnstaedt SD, Gottwein E, Skalsky RL, Luftig MA, Cullen BR. Virally induced cellular microRNA miR-155 plays a key role in B-cell immortalization by Epstein-Barr virus. J Virol. 2010;84(22):11670–8.

32. Kurata JS, Lin RJ. MicroRNA-focused CRISPR-Cas9 library screen reveals fitness-associated miRNAs. RNA. 2018;24(7):966–81.

33. Colic M, Wang G, Zimmermann M, Mascall K, McLaughlin M, Bertolet L, et al. Identifying chemogenetic interactions from CRISPR screens with drugZ. Genome Med. 2019;11(1):52.

34. Trissal MC, Wong TN, Yao JC, Ramaswamy R, Kuo I, Baty J, et al. MIR142 Loss-of-Function Mutations Derepress ASH1L to Increase HOXA Gene Expression and Promote Leukemogenesis. Cancer Res. 2018;78(13):3510–21.

35. Urbanek-Trzeciak MO, Galka-Marciniak P, Nawrocka PM, Kowal E, Szwec S, Giefing M, et al. Pan-cancer analysis of somatic mutations in miRNA genes. EBioMedicine. 2020;61:103051.

36. Kwanhian W, Lenze D, Alles J, Motsch N, Barth S, Doll C, et al. MicroRNA-142 is mutated in about 20% of diffuse large B-cell lymphoma. Cancer Med. 2012;1(2):141–55.

37. Kramer NJ, Wang WL, Reyes EY, Kumar B, Chen CC, Ramakrishna C, et al. Altered lymphopoiesis and immunodeficiency in miR-142 null mice. Blood. 2015;125(24):3720–30.

38. Motsch N, Alles J, Imig J, Zhu J, Barth S, Reineke T, et al. MicroRNA profiling of Epstein- Barr virus-associated NK/T-cell lymphomas by deep sequencing. PLoS One. 2012;7(8):e42193.

39. Ambrosio MR, Navari M, Di Lisio L, Leon EA, Onnis A, Gazaneo S, et al. The Epstein Barr- encoded BART-6-3p microRNA affects regulation of cell growth and immuno response in Burkitt lymphoma. Infect Agent Cancer. 2014;9:12.

40. Wu H, Ye C, Ramirez D, Manjunath N. Alternative processing of primary microRNA transcripts by Drosha generates 5’ end variation of mature microRNA. PLoS One. 2009;4(10):e7566.

41. Manzano M, Forte E, Raja AN, Schipma MJ, Gottwein E. Divergent target recognition by coexpressed 5’-isomiRs of miR-142-3p and selective viral mimicry. RNA. 2015;21(9):1606–20.

42. Fachko DN, Chen Y, Skalsky RL. Epstein-Barr Virus miR-BHRF1-3 Targets the BZLF1 3’UTR and Regulates the Lytic Cycle. J Virol. 2022;96(4):e0149521.

43. Corcoran DL, Georgiev S, Mukherjee N, Gottwein E, Skalsky RL, Keene JD, et al. PARalyzer: definition of RNA binding sites from PAR-CLIP short-read sequence data. Genome Biol. 2011;12(8):R79.

44. Majoros WH, Lekprasert P, Mukherjee N, Skalsky RL, Corcoran DL, Cullen BR, et al. MicroRNA target site identification by integrating sequence and binding information. Nat Methods. 2013;10(7):630–3.

45. Ma Z, Liu T, Huang W, Liu H, Zhang HM, Li Q, et al. MicroRNA regulatory pathway analysis identifies miR-142-5p as a negative regulator of TGF-beta pathway via targeting SMAD3. Oncotarget. 2016;7(44):71504–13.

46. Berrien-Elliott MM, Sun Y, Neal C, Ireland A, Trissal MC, Sullivan RP, et al. MicroRNA-142 Is Critical for the Homeostasis and Function of Type 1 Innate Lymphoid Cells. Immunity. 2019;51(3):479–90 e6.

47. Sun Y, Oravecz-Wilson K, Bridges S, McEachin R, Wu J, Kim SH, et al. miR-142 controls metabolic reprogramming that regulates dendritic cell activation. J Clin Invest. 2019;129(5):2029–42.

48. Ge SX, Jung D, Yao R. ShinyGO: a graphical gene-set enrichment tool for animals and plants. Bioinformatics. 2020;36(8):2628–9.

49. Wu L, Cai C, Wang X, Liu M, Li X, Tang H. MicroRNA-142-3p, a new regulator of RAC1, suppresses the migration and invasion of hepatocellular carcinoma cells. FEBS Lett. 2011;585(9):1322–30.

50. Yasuda T, Kometani K, Takahashi N, Imai Y, Aiba Y, Kurosaki T. ERKs induce expression of the transcriptional repressor Blimp-1 and subsequent plasma cell differentiation. Sci Signal. 2011;4(169):ra25.

51. Matallanas D, Birtwistle M, Romano D, Zebisch A, Rauch J, von Kriegsheim A, et al. Raf family kinases: old dogs have learned new tricks. Genes Cancer. 2011;2(3):232–60.

52. Hodge C, Liao J, Stofega M, Guan K, Carter-Su C, Schwartz J. Growth hormone stimulates phosphorylation and activation of elk-1 and expression of c-fos, egr-1, and junB through activation of extracellular signal-regulated kinases 1 and 2. J Biol Chem. 1998;273(47):31327–36.

53. Larkin J, Ascierto PA, Dreno B, Atkinson V, Liszkay G, Maio M, et al. Combined vemurafenib and cobimetinib in BRAF-mutated melanoma. N Engl J Med. 2014;371(20):1867–76.

54. Dean L, Kane M. Dabrafenib Therapy and BRAF Genotype. In: Pratt VM, Scott SA, Pirmohamed M, Esquivel B, Kattman BL, Malheiro AJ, editors. Medical Genetics Summaries. Bethesda (MD) 2012.

55. Gureasko J, Galush WJ, Boykevisch S, Sondermann H, Bar-Sagi D, Groves JT, et al. Membrane-dependent signal integration by the Ras activator Son of sevenless. Nat Struct Mol Biol. 2008;15(5):452–61.

56. Gorbea C, Mosbruger T, Cazalla D. A viral Sm-class RNA base-pairs with mRNAs and recruits microRNAs to inhibit apoptosis. Nature. 2017;550(7675):275-9.

57. Riley KJ, Rabinowitz GS, Yario TA, Luna JM, Darnell RB, Steitz JA. EBV and human microRNAs co-target oncogenic and apoptotic viral and human genes during latency. EMBO J. 2012;31(9):2207–21.

58. Skalsky RL, Corcoran DL, Gottwein E, Frank CL, Kang D, Hafner M, et al. The viral and cellular microRNA targetome in lymphoblastoid cell lines. PLoS Pathog. 2012;8(1):e1002484.

59. Skalsky RL, Barr SA, Jeffery AJ, Blair T, Estep R, Wong SW. Japanese Macaque Rhadinovirus Encodes a Viral MicroRNA Mimic of the miR-17 Family. J Virol. 2016;90(20):9350–63.

60. Gottwein E, Corcoran DL, Mukherjee N, Skalsky RL, Hafner M, Nusbaum JD, et al. Viral microRNA targetome of KSHV-infected primary effusion lymphoma cell lines. Cell Host Microbe. 2011;10(5):515–26.

61. Trobaugh DW, Gardner CL, Sun C, Haddow AD, Wang E, Chapnik E, et al. RNA viruses can hijack vertebrate microRNAs to suppress innate immunity. Nature. 2014;506(7487):245-8.

62. Hart T, Tong AHY, Chan K, Van Leeuwen J, Seetharaman A, Aregger M, et al. Evaluation and Design of Genome-Wide CRISPR/SpCas9 Knockout Screens. G3 (Bethesda). 2017;7(8):2719- 27.

63. Martin M. Cutadapt removes adapter sequences from high-throughput sequencing reads. 2011. 2011;17(1):3.

64. Langmead B, Salzberg SL. Fast gapped-read alignment with Bowtie 2. Nat Methods. 2012;9(4):357–9.

65. Li H, Handsaker B, Wysoker A, Fennell T, Ruan J, Homer N, et al. The Sequence Alignment/Map format and SAMtools. Bioinformatics. 2009;25(16):2078–9.

66. Langmead B, Trapnell C, Pop M, Salzberg SL. Ultrafast and memory-efficient alignment of short DNA sequences to the human genome. Genome Biol. 2009;10(3):R25.

